# Cosmic silence and viral noise: transcriptomic crosstalk in *Caenorhabditis elegans* under simulated space conditions

**DOI:** 10.64898/2025.12.02.691756

**Authors:** Ana Villena-Giménez, Esmeralda G. Legarda, Rubén González, Victoria G. Castiglioni, Santiago F. Elena

## Abstract

Spaceflight environments pose unique physiological challenges due to altered gravity and radiation exposure. To investigate how these abiotic stressors interact with viral infection, we analyzed the transcriptomic response of *Caenorhabditis elegans* acclimated to simulated microgravity (µG) and below-background muon radiation flux (BBR), upon infection with Orsay virus (OrV). Using RNA-seq, we characterized gene expression profiles across single and combined stress conditions. Both µG and BBR elicited distinct stress responses, including modulation of oxidative stress, lipid metabolism, and immune pathways. OrV infection alone induced robust transcriptional changes, but its impact was significantly attenuated when combined with either abiotic stress, suggesting antagonistic interactions. Notably, proviral genes such as *drl-1*, *fat-7* and *hipr-1* were downregulated under BBR and µG, potentially impairing viral replication. Gene ontology analyses revealed enrichment in immune effectors, RNA metabolism, and proteostasis-related pathways, particularly under BBR. Viral load and RNA2/RNA1 ratios were reduced in both stress conditions, indicating a shift in viral replication dynamics. Moreover, genomic diversity and defective viral genome formation were differentially affected, with increased genetic diversity and structural variation under stress. These findings suggest that acclimation to off-Earth conditions primes the host for a dampened response to an acute viral infection, potentially through resource reallocation and transcriptional attenuation. This study provides transcriptomic insight into viral infection under space-relevant conditions, highlighting complex stress interactions and their implications for host-pathogen dynamics in extraterrestrial environments.

## 1. Introduction

Space exploration is accelerating, with new missions to the Moon and Mars poised to expose organisms to combined stressors, including altered gravity and non-terrestrial radiation fields. Microgravity (∼ 0 g) disrupts human physiology across multiple systems, notably musculoskeletal and cardiovascular function (Tanaka et al. 2017). The immune system is also affected: astronauts exhibit altered lymphocyte distributions (Grove et al. 1995), reduced NK cell activity (Konstantinova et al. 1995), elevated proinflammatory mediators such as IL-6 and cortisol (Stein & Schluter 1994), and depressed virus-specific T-cell responses (Crucian et al. 2013). Ground-based microgravity (µG) simulations recapitulate increased inflammatory tone and reduced T-and NK-cell function (Wu et al. 2024). Clinically relevant outcomes include viral reactivation during missions: Epstein-Barr virus (EBV), varicella-zoster virus, and cytomegalovirus can increase in frequency, duration, and copy number even during short flights (Mehta et al. 2017). In mice flown to space, transcriptomics reveals up-regulation of virus-related pathways (*e.g*., prion disease, coronavirus disease, hepatitis A, B and C) (Zhang et al. 2024). Yet, findings under simulated µG are mixed: Kaposi’s sarcoma-associated herpesvirus (Honda et al. 2020) and EBV (Long et al. 1999) can remain latent, whereas latent retroviral transcription can be induced in human immune cells after ∼ 25 h of µG (Wu et al. 2024). Microbiome-associated viromes also appear dynamic: phages and other potential viral pathogens within the human microbiota increase activity during in-flight phases (Tierney et al. 2024). Thus, the net impact of µG on viral dynamics remains unresolved.

Radiation exposure is a parallel challenge. On the International Space Station (ISS), annual effective doses (∼ 110 - 180 mSv year^−1^) exceed those on Earth (∼ 0.5 - 70 mSv year^−1^) (Restier-Verlet et al. 2021). Crucially, radiation composition differs: space exposures include galactic cosmic rays, solar particle events, and trapped belt electrons/protons (Freese et al. 2016), while terrestrial exposure is dominated by telluric sources (*e.g*., radon), radioactive materials, and attenuated cosmic rays (Restier-Verlet et al. 2021). On Earth’s surface, ∼85% of detected cosmic radiation comprises muons, *i.e*., low-linear energy transfer (LET), highly penetrating particles with high flux (Atri & Melott 2011, Atri & Melott 2014). These muons are largely absent in deep underground environments; intriguingly, below-background radiation (BBR) studies in such facilities report increased radiosensitivity (Carbone et al. 2009), diminished cellular defense mechanisms (Fratini et al. 2015), and broad transcriptional reprogramming: down-regulated primary metabolism with up-regulated immune and stress-response pathways (Zarubin et al. 2021). These observations have led to the hypothesis that chronic muon flux contributes to normal physiological set points, and that BBR can perturb cellular homeostasis.

The nematode *Caenorhabditis elegans* is a proven workhorse for space biology due to ease of culture, rapid life cycle, compact size, and deep molecular annotation with substantial human gene homology (Lai et al. 2000). The discovery of the *Nodavirus*-like Orsay virus (OrV), a mild, nonlethal, intestinal pathogen transmitted orofecally, enables host-virus studies in a whole-animal, genetically tractable system (Félix et al. 2011). In *C. elegans*, biotic and abiotic stressors can interact nonlinearly: for example, ZnO nanoparticles or *Klebsiella pneumoniae* alone suppress reproduction, yet together show no effect (Cochran et al. 2023). Mild stress preconditioning can increase survival under subsequent severe stress, including diet-dependent cadmium resilience (Dölling et al. 2019). Crosstalk between abiotic stress and infection is bidirectional: OrV infection increases heat-shock resistance (Castiglioni & Elena 2024), while heat stress increases resistance to OrV (Huang et al. 2021), with overlapping transcriptional responses enriched for *pals*-family genes and intracellular pathogen response modules (Zhou et al. 2019; Huang et al. 2021). OrV infection during larval development progresses through four distinct phases: prereplication (0 - 6 hpi), exponential replication peaking at 12 hpi, moderate replication (14 - 36 hpi), and persistent residual infection (38 - 44 hpi) (Castiglioni, Olmo-Uceda, Villena-Giménez et al. 2024). After the acute phase, a latent infection is stablished (Castiglioni et al. 2025).

We previously described complex interactions among OrV infection, simulated µG, and BBR (Villena-Giménez et al. 2025). In this previous study, we employed a fully factorial experimental design examining how µG, BBR, and OrV infection, alone and in all combinations, influences physiological traits and viral load. While BBR radically affected viral accumulation dynamics, µG had a minor effect. Both factors significantly impacted reproduction and morphology, with some effects magnified by viral infection. These results revealed how even partial modifications of Earth-like gravity and radiation levels can alter pathogen-host interactions. Other studies, have shown that simulated µG recapitulates spaceflight-like gene expression shifts in *C. elegans*, especially in cytoskeletal/muscle and metabolic pathways (Higashibata et al. 2016), and a core of ∼118 µG-responsive genes enriched for locomotion, morphogenesis, and cuticle biology —44 of which map to 64 human orthologs (Çelen et al. 2023). In BBR, *C. elegans* transcriptomes show up-regulated sperm proteins and down-regulated cuticle/collagen genes (Van Voorhies et al. 2020). However, the interplay between viral infection and the combined stresses of µG and reduced muon flux remains unexplored.

Here, we characterize the transcriptomic landscape of OrV infection in *C. elegans* acclimated to simulated space conditions, building on Villena-Giménez et al. (2025). We exposed larvae populations to simulated µG, BBR, and OrV infection, each alone and in combination (µG + OrV and BBR+ OrV), following two-generation acclimation to the abiotic stressors. From the virus standpoint, we quantify how these abiotic stresses shape OrV accumulation and intra-host diversity.

## 2. Materials and methods

### 2.1. C. elegans strains and culturing

*C. elegans* was cultured and maintained at 20 °C on nematode growth media (NGM) agar plates seeded with *Escherichia coli* OP50. ERT54 (*jyIs8[pals-5p::GFP + myo-2p::mCherry]X*), a transgenic strain with a genetic wild-type (Bristol N2) background that expresses GFP in response to intracellular infection (Bakowski et al. 2014) was used for all experiments. The more susceptible strain SFE2 (*drh-1(ok3495)IV;mjls228*) was used to produce OrV stocks.

In order to obtain synchronized animal populations, plates with embryos were carefully washed with M9 buffer to remove larvae and adults but leaving the embryos behind. Plates were washed again using M9 buffer after 1 h to collect larvae hatched within that time span and transferred to seeded NGM plates.

### 2.2. Microgravity simulation

Experiments were performed as previously described (Villena-Giménez et al. 2025). Briefly, a random position machine (RPM) (Yuri Gravity GmbH) was employed to simulate microgravity conditions selecting the zero-gravity mode. Values of *g*-force were monitored through the RPM software and maintained at approximately 0.001 - 0.002 g throughout the experiments. The RPM was placed inside an incubator at 20 °C.

Animals used in experiments of µG conditions were acclimated along two generations, with studies done on the third generation and its progeny. Plates were sealed with parafilm to maintain their humidity.

### 2.3. Below-background radiation conditions

Experiments were performed in the LAB2400 at the Canfranc Underground Laboratory (LSC), located in Estación de Canfranc (Huesca, Spain) as previously described (Villena-Giménez et al. 2025). The measured integrated muon radiation flux in this facility is ∼ 0.005 m^−2^ s^−1^ (Trzaska et al. 2019). For comparison, the muon radiation flux at sea level in the Northern hemisphere is about 150 m^−2^ s^−1^ (Particle Data Group et al. 2022).

Thermoluminescent dosimeters sensitive to various sources of radiation are placed at different locations of the underground laboratory. In 2022 they showed an average dose rate of 0.71 ±0.03 mSv year^−1^, compared with the 1.36 ±0.03 mSv year^−1^ of the above ground laboratory (Hernández-Antolín et al. 2024). About radiation produced by radon, a good ventilation system and a Radon Abatement System got to lower radiation levels to 0.001 Bq m^−3^, substantially lower levels compared with the 200 - 280 Bq m^−3^ measured at the surface laboratory (Pérez-Pérez et al. 2022).

Nematodes used in BBR experiments were acclimated to this BBR condition for two generations prior to experiments.

### 2.4. Viral stock preparation, virus quantification and inoculation procedure

For OrV (strain JUv1580_vlc) stock preparation, SFE2 animals were inoculated as previously described (Castiglioni, Olmo-Uceda, Martín et al. 2024). In short, animals were allowed to grow for 5 days and then resuspended in M9 (0.22 M KH_2_PO_4_, 0.42 M Na_2_HPO_4_, 0.85 M NaCl, 1 mM MgSO_4_), let stand for 15 min at room temperature, vortexed, and centrifuged for 2 min at 400 g. The supernatant was centrifuged twice at 21,000 g for 5 min and then passed through a 0.2 μm filter. RNA of the resulting viral stock was extracted using the Viral RNA Isolation kit (NZYTech). The concentration of viral RNA was then determined by RT-qPCR using a standard curve, and normalized across different stocks (details below). Primers used for RT-qPCRs can be found in Table S1.

For the standard curve, cDNA of JUv1580_vlc was obtained using AccuScript High-Fidelity Reverse Transcriptase (Agilent) and reverse primers at the 3’ end of the genome. Approximately 1000 bp of the 3’ end of RNA2 were amplified using forward primers containing 20 bp coding the T7 promoter and DreamTaq DNA Polymerase (Thermo Fisher). The PCR products were gel purified using MSB Spin PCRapace (Invitek Molecular) and an *in vitro* transcription was performed using T7 Polymerase (Merck). The remaining DNA was then degraded using DNase I (Life Technologies). RNA concentration was determined by NanoDrop (Thermo Fisher) and the number of molecules per µL was determined using the online tool EndMemo RNA Copy Number Calculator (https://www.endmemo.com/bio/dnacopynum.php). Primers used for the standard curve can be found in Table S1.

For inoculation experiments, synchronized populations were inoculated by pipetting 60 μL of viral stock on top of the bacterial lawn containing the animals. The normalized inoculum contained 2.6×10^7^ copies of OrV RNA2/μL. The efficiency of this viral stock (measured as animals showing activation of the *pals-5p::GFP* reporter at 48 hpi) was 72 ±3 % (mean ±1 SEM, *n* = 5 plates with 44 - 48 animals per plate).

### 2.5. RNA extractions

Sample preparation and RNA extractions were performed as previously described (Castiglioni, Olmo-Uceda, Villena-Giménez et al. 2024). Synchronized populations of 300 inoculated and control animals were collected at 14 hpi with PBS-0.05% Tween. Samples of inoculated animals were performed in triplicates. Samples were centrifuged for 2 min at 1350 rpm and the supernatant was discarded. Another two wash steps were performed before freezing the samples in liquid nitrogen. 500 µL of Trizol (Invitrogen) were added to the nematode pellet and disrupted by following five cycles of freeze-thawing and five cycles of 30 seconds of vortex followed by 30 seconds of rest. 100 µL of chloroform were then added and the tubes were shaken for 15 s and let rest for 2 min. Samples were centrifuged for 15 min at 11,000 g at 4 °C and the top layer containing the RNA was then mixed with the same volume of 100% ethanol. The sample was then loaded into RNA Clean & Concentrator columns (Zymo Research) and the rest of the protocol was followed according to manufacturer instructions.

### 2.6. Total RNA extraction and preparation for RNA-seq

Sample preparation was performed as previously described (Castiglioni, Olmo-Uceda, Villena-Giménez et al. 2024). Library preparation and Illumina sequencing was done by Novogene Europe (www.novogene.com) using a NovaSeq 6000 platform and a lnc-stranded mRNA-seq library method, ribosomal RNA depletion, directional library preparation, 150 paired end, and 6-Gb raw data per sample. Novogene checked the quality of the libraries using a Qubit 4 Fluorometer (Thermo Fisher Scientific), qPCR for quantification, and Bioanalyzer for size distribution detection.

### 2.7. RNA-seq and host data processing

The quality of the resulting fastq files was checked using FastQC (Andrews 2010), MultiQC (Ewels et al. 2016). A preprocess step running bbduk.sh was included to remove adapters and trim read ends with low quality (trimq = 10). Then, reads were mapped against the reference genome of *C. elegans* by STAR (Dobin et al. 2013), giving as input the genome of N2 Bristol and its annotations (GCF_000002985.6_WBcel235), downloaded from NCBI.

To obtain the matrix of counts per gene and sample, the function summarizeOverlaps from the package GenomicAlignments (Lawrence et al. 2013) was employed (strand-aware, for pair-end reads and mode = “Union”). This matrix count was annotated using the dataset wbps_gene from parasite_mart, AnnotationDbi and org.Ce.eg.db. Exploratory methods such as principal components analysis (PCA) and clustering in R were visualized, with a previous variance stabilizing transformation (VST).

We used DESeq2 (Love et al. 2014) for the differential expression analysis, applying a filter of minimum of ten counts in at least three samples (size of our groups). Previously, factors of unwanted variation (*W*_1_ + *W*_2_) using replicate samples were calculated with the R package RUVSeq (Risso et al. 2014). The complete design was set as ∼ *W*_1_ + *W*_2_ + virus + treatment + virus×treatment and using Wald test. A likelihood-ratio test (LRT) was used to confirm the need of taking into account the interaction factor. IHW package was used for the multiple testing procedure and DESeq2 to shrink log_2_ fold changes (*FC*). DEGs were considered significant with Benjamini and Hochberg adjusted *P* < 0.05 values. In heatmaps, asterisks indicate significance level.

### 2.8. Data visualization

Venn diagrams were generated using the R library VennDiagram, including for each group only genes with adjusted *P* < 0.05. The same number of genes were given as input to the following functional analyses. The over-representation analysis (ORA) with the hyperGTest method of Gostats (Falcon & Gentleman 2007) was run using a cutoff *P* = 0.05 and all genes from org.Ce.eg.db as universe.

Gene networks were created with Cytoscape v3.10.4 (Shannon et al. 2003) selecting the option “Full STRING network” and a confidence cutoff of 0.4 and no additional interactors. For community clustering, GLay from clusterMaker2 (Morris et al. 2011) was employed.

### 2.9. Virus characterization

After the pre-processing steps of fastq files, bwa-mem (Li 2013) was employed for mapping OrV reads. Normalized quantification of viral load was calculated dividing the viral counts by the host counts. For sequence diversity, we chose normalized Shannon entropy (*H_n_*) per position, calculated as in (Gregori et al. 2014). We also used the number of positions of each sample that were distributed in the top 5% of all entropy values. For non-standard viral genomes (nsVG) inspection, DVGfinder (Olmo-Uceda et al. 2022) was run using the metasearch mode for the two segments of OrV genome separately, after sequences mapped against host genome were removed. We applied stringent filtering strategies to the results, retaining only nsVGs with a minimum length of five nucleotides and supported by at least ten independent reads by one or two of the algorithms implemented, ViReMa (Sotcheff et al. 2023) and DI-tector (Beauclair et al. 2018), thereby restricting the analysis to high-confidence events. We did not segregate the data by the sense of the nsVGs. The abundance of nsVGs was normalized and expressed in terms of counts per 100,000 viral reads (viral counts per hundred thousand reads; VCPHT).

## 3. Results and discussion

### 3.1. Stress acclimation produces antagonistic transcriptional interactions with viral infection

In this study, we investigated how off-Earth conditions interact with viral infection at the gene expression level. To achieve this, we used a random positioning machine (RPM) to simulate microgravity (µG) and conducted experiments in the Canfranc Underground Laboratory (LSC) to reduce muon flux radiation below natural background levels (BBR). These experiments were performed in the *C. elegans* - OrV pathosystem. Importantly, we aimed to capture long-term stress responses to these abiotic stresses rather than acute reactions, so animals were acclimated for two generations to the abiotic stressors (µG or BBR), given that intergenerational stress effects can persist for up to three generations (Kishimoto et al. 2017). Nematodes were inoculated immediately after hatching with a high-concentration OrV inoculum and collected at 14 hpi, corresponding to the peak viral load previously reported (Castiglioni et al., 2024; Villena-Giménez et al. 2025).

We first quantified the infection effect on the host transcriptional response within each background (Fig. 1A, Fig. S1) using a general linear model (∼ virus + treatment + treatment × virus). Across conditions, OrV elicited a robust core response (67 upregulated, seven downregulated genes; Fig. 1B - 1C, Fig. S2A-S2B, Table S2), indicating a conserved infection program that persists despite environmental acclimation.

**Fig. 1.**
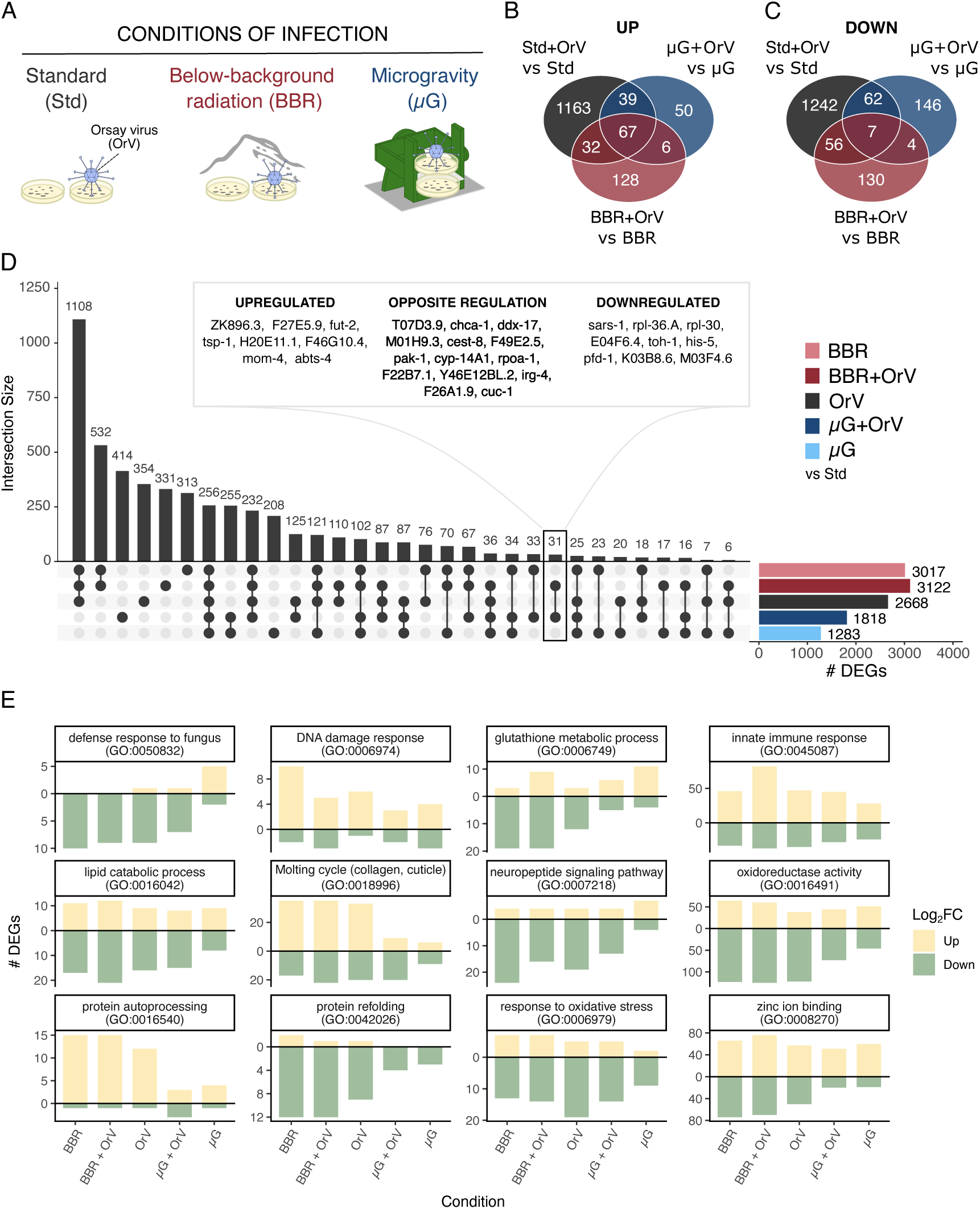
(A) Schematic representation of the experimental design. (B) Venn diagram showing upregulated and downregulated genes across infection conditions, based on contrasts between infected and non-infected samples within each condition. (C) Upset plot illustrating exclusive set intersections among physical and infection conditions compared to non-infected nematodes under standard conditions. The total size of each set is shown in the right bar plot, while all possible intersections are represented below, with their frequencies displayed in the top bar plot. (D–E) Comparison of infection conditions by GO categories, showing the number of differentially expressed genes associated with each term, using the same contrast strategy as in (C).

To benchmark stress magnitude, we contrasted each treatment to non-infected nematodes under standard conditions (Fig. 1D, Fig. S1G-S1H). BBR and OrV alone produced similarly large transcriptomic responses (3,017 and 2,668 DEGs; adjusted *P* < 0.05), whereas µG yielded fewer (1,283). When combined with infection, the overall response intensified under BBR (3,122 DEGs) but was partially antagonistic under µG (1,818). Interaction terms were substantial: 1,940 DEGs for BBR × virus and 1,159 for µG × virus, with 947 shared between interactions (Fig. S2). These patterns reveal a conserved coupling among stress pathways. GO summaries further highlighted recurring modulation of proteostasis (*e.g*., protein auto processing and refolding), oxidative stress/oxidoreductase activity, lipid catabolism, and zinc binding, consistent with infection-driven proteotoxic load (Chen et al., 2017; Reddy et al., 2017), the lipid and zinc dependence of replication (Casorla-Perez et al., 2022), and spaceflight-linked lipid remodeling (Adenle et al., 2009) (Fig. 1E). Notably, OrV alone elicited the broadest oxidative-stress response, which was blunted by prior acclimation to BBR or µG, consistent with stress conditioning.

### 3.2. Proviral and lipid metabolism genes are suppressed by BBR and µG

We next examined how BBR and µG affect expression of proviral genes, host genes required for OrV infection (Table S3). BBR modestly upregulated *alg-1* (Fig. S2D; log₂*FC* = 0.573, adjusted *P* = 0.001) and downregulated *drl-1* (Fig. S2D; log_2_*FC* = −0.788, adjusted *P* = 0.007) and *hipr-1* (Fig. S2D; log_2_*FC* = −0.379, adjusted *P* = 0.005), three proviral factors essential during early OrV replication (Parker et al., 2007; Sandoval et al., 2019; Jiang et al., 2020). In parallel, OrV infection suppressed fatty-acid regulators (*elo-1*, *elo-2*, *fat-6*, *fat-7*, *nhr-49*, and *nhr-80*) at 14 hpi (Fig. S2E), a pattern recapitulated under BBR. In the comparison BBR + OrV *vs* BBR, only *fat-7* exhibited a significant reduction (log_2_*FC* = −0.821, adjusted *P* = 0.002) and, moreover, interaction terms were significant across this lipid module, indicating non-additive control and may pinpointing the existence of a regulatory limit. Under µG, *fat-6*, *fat-7* and *nhr-80* were also downregulated (all adjusted *P* ≤ 0.009), with no significant regulation in the comparison µG + OrV *vs* µG, and significant µG × OrV interactions for *elo-1*, *fat-7*, *nhr-49*, and *nhr-80* (Fig. S2E). Together, these results position lipid metabolism as a shared lever through which µG and BBR shape early infection dynamics and host fitness traits. This observation aligns with the broad roles of lipids in viral entry and replication (Heaton & Randall, 2011) and the requirement of *sbp-1* for OrV replication (Casorla-Perez et al., 2022).

### 3.3. BBR and µG attenuate canonical infection responses

STRING-based clustering of infection-responsive genes uncovered modules whose amplitude depended on the physical background, showing stronger differences in µG+OrV network (Fig. 2A). To assess whether antagonistic interactions affect specific infection pathways, we examined expression of previously characterized OrV infection-responsive genes (Castiglioni et al., 2024), whose cluster exhibits most robust expression (Fig. 2A-2B). Under BBR alone, *B0507.8*, *dod-22* and *eol-1* were mildly increased, while *C45B2.2*, *gst-33* and *ZK792.4* were reduced. In BBR + OrV, 11 genes, including IPR/innate-immunity effectors *pals-5*, *pals-6*, *pals-14*, *pals-27*, and *F26F2.1*, showed weaker upregulation than under OrV alone (Huang et al., 2021; Lažetić et al., 2022; Castiglioni et al., 2024). The infection-inducible *eol-1* (Shen et al., 2014) and *trpl-5* (Goodman, 2006) also attenuated, indicating partial priming/antagonism. Conversely, *ZK792.4* and *C45B2.2* downregulation under BBR was not significantly altered by OrV, maintaining BBR-baseline levels.

**Fig. 2.**
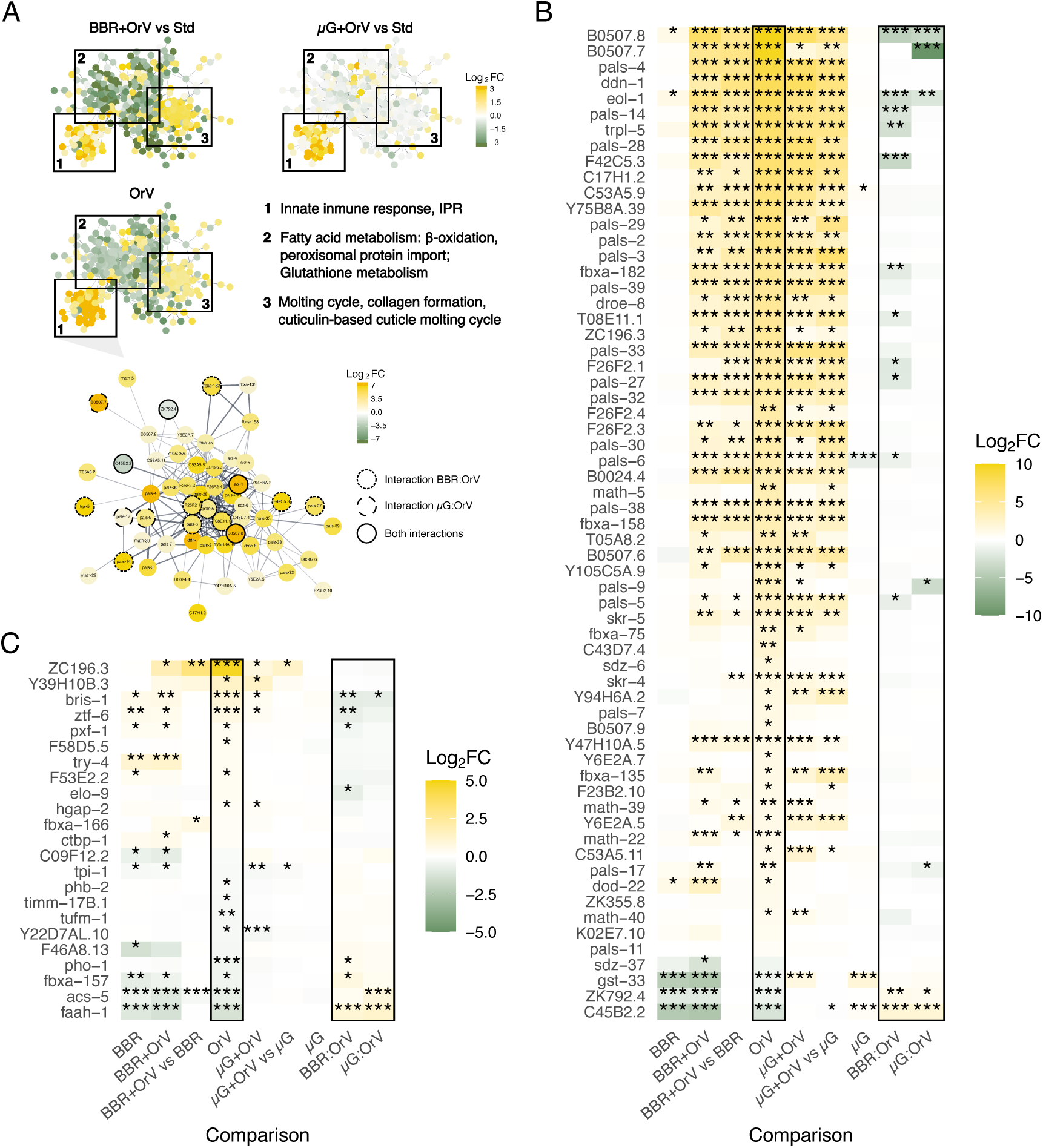
(A) Cluster from a STRING-based network of genes with ∣log_2_*FC*∣ > 1 in response to OrV infection, including infection-specific response genes. The three main gene communities are highlighted with squares, and their predominant biological processes are indicated. In the community of infection-specific response genes, log_2_*FC* values are shown for both BBR + OrV *vs* standard (Std) and µG + OrV *vs* Std comparisons. In the community of infection-specific response genes, circle color also represents log₂*FC* values, while the outline style denotes genes showing significant interaction effects between BBR or µG and OrV infection. Asterisks denote adjusted significance levels (\**P* < 0.05, \*\**P* < 0.01, \*\*\**P* < 0.001). (B) Genes from panel A showing log_2_*FC* across different conditions (relative to non-infected nematodes in each condition, unless otherwise indicated) and for the interactions BBR × OrV and µG × OrV. (C) Genes previously identified as correlating or anti-correlating with OrV accumulation, displaying log_2_*FC* across conditions and for the interactions BBR × OrV and µG × OrV.

For µG + OrV, five IPR components (*B0507.7*, *B0507.8*, *eol-1*, *pals-9*, and *pals-17*) displayed lower expression than the additive expectation from the single stresses (Lažetić et al., 2022; Lažetić et al., 2023). Load-correlated infection markers (*bris-1*, *elo-9*, *pxf-1*, and *ztf-6*) were buffered by µG and BBR, and both stresses interfered with the canonical downregulation of *faah-1* (Nandakumar & Tan, 2008; Harrison et al., 2014; Ruiz et al., 2019) (Fig. 2C). Altogether, these trends support a stress-primed, antagonistic interaction: the environmental background tempers hallmark infection programs while preserving selective defense and metabolic adjustments.

### 3.4. Stress-specific transcriptional signatures reveal distinct strategies in response to viral infection

Beyond shared antagonism, BBR and µG produced stress-specific transcriptional signatures. Under BBR + OrV (*vs* BBR), we detected 233 upregulated and 197 downregulated genes, largely overlapping with the standard infection signature. Excluding genes shared with individual stresses yielded 89 and 94 upregulated and downregulated genes, respectively, what was reduced to 15 uniquely regulated genes by filtering for |log_2_*FC*| > 1 (Fig. 3A). These included *sri-36* (serpentine receptor), tetraspanins *tsp-1* and *tsp-2*, and ion-binding proteins such as *Y69A2AL.2* (phospholipid-binding prediction), consistent with barrier remodeling and durable stress memory (Jiang et al., 2024). Conversely, mtl-2 and several cytochrome P450s (*e.g*., *cyp-35A3, cyp-35A5* and *cyp-35C1*) decreased, alongside *clec-206*, suggesting deliberate restraint of detoxification to limit reactive intermediates (Hall et al., 2012). GO terms enriched among upregulated genes included immune defense, and notably, the predominant PERK-mediated UPR (Kim et al., 2020; Brown et al., 2021; Zheng et al., 2022; Son et al., 2024; Castiglioni et al., 2024), reinforcing that BBR acts as a physiological stressor and aligning with hormesis-like responses. GO terms overrepresented in downregulated genes comprises rRNA metabolism and ribosome biogenesis, consistent with proteostasis challenges.

**Fig. 3.**
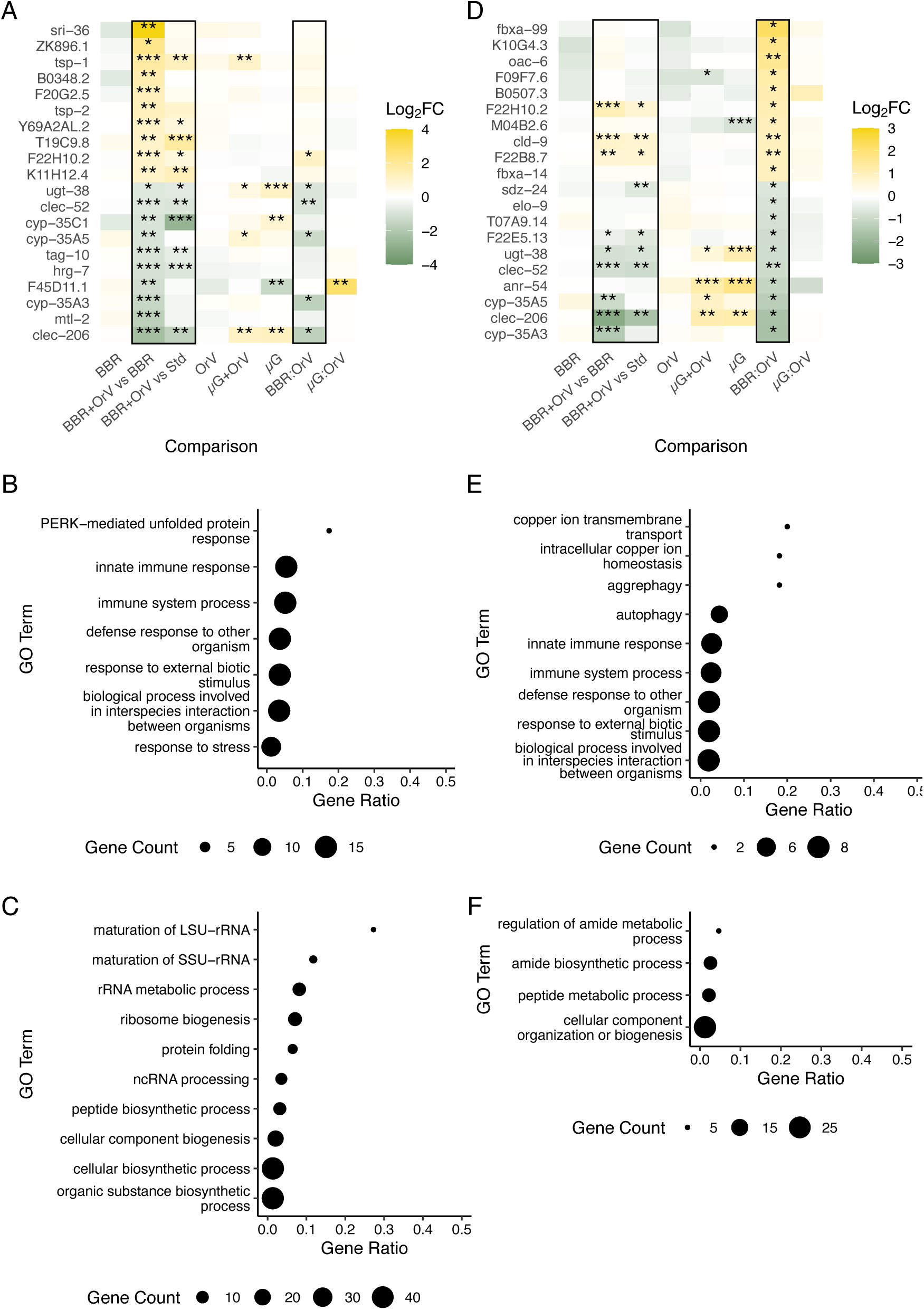
(A) Panel of most significantly upregulated or downregulated genes exclusively under BBR conditions after OrV inoculation (14 hpi), compared to non-infected nematodes in this condition. Log_2_*FC* values across all analyzed conditions and interaction effects are shown. Unless otherwise stated, all comparisons were made relative to non-infected nematodes maintained under standard conditions (Std). Asterisks indicate adjusted significance levels (\**P* < 0.05, \*\**P* < 0.01, \*\*\**P* < 0.001). (B) GO enrichment analysis of top biological processes based on exclusively and significantly (adjusted *P* < 0.01) upregulated genes from the comparison in (A), presented as in (B). (C) GO enrichment analysis of top biological processes based on exclusively and significantly (adjusted *P* < 0.01) downregulated genes from the comparison in (A). (D) Panel of the ten most significantly upregulated and downregulated genes exclusively for the BBR × OrV interaction, displayed as in (A). (E) GO enrichment analysis of top biological processes based on exclusively and significantly upregulated genes for this interaction, presented as in (B). (F) GO enrichment analysis of top biological processes based on exclusively and significantly downregulated genes for this interaction, presented as in (B).

Among genes showing significant interaction between BBR and infection status, we identified 1,940 genes; among those showing the strongest interaction, we found *argk-1* (creatine kinase linked to stress resistance/lifespan), *col-147* (collagen), *fipr-7* (feeding), *ttr-21*, and *Y73F4.2* (pathogen response) (McQuary et al., 2016; Burton et al., 2021; Sheshadri et al., 2021; Li et al., 2025). In contrast, infection-inducible genes *B0507.8*, *eol-1*, *F42C5.3* were attenuated (higher than BBR alone, lower than OrV alone) (Teuscher et al., 2019; van Sluijs et al., 2022; Castiglioni et al., 2024) (Fig. S3A). GO analysis with |log_2_*FC*| > 1 genes showed supra-additive enrichment for lipid catabolism, detoxification, and immunity among interaction-up genes (383), while interaction-down genes (262) emphasized cuticle development, genitalia, sensory development, and protein auto processing (Fig. S3B). Collagen modules, linked to BBR environments and antiviral activity, were prominent (Van Voorhies et al., 2020; Zhou et al., 2024), consistent with prior observations of altered reproduction under BBR + OrV relative to BBR alone (Villena-Giménez et al., 2025). Excluding significant genes shared with BBR or OrV effects, 59 and 66 genes resulted to be interaction-specific inducted and repressed, respectively (Fig. 3D). GO enrichment revealed that the upregulated exclusive genes were related to copper homeostasis, autophagy and immune system (Fig. 3E), while the downregulated showed amide biosynthesis and peptide metabolism enriched under combined stress conditions (Fig. 3F).

OrV infection upon microgravity (µG + OrV *vs* µG) produced fewer unique genes, eight with ∣log₂*FC*∣ > 1, out of which 48 upregulated and 71 downregulated genes were absent from single-stress lists, including *pals-31* and *chil-19* (carbohydrate metabolism) (Fig. 4A). GO terms amongst these downregulated genes pointed to amide biosynthesis, peptide metabolism, and metal transport and homeostasis (particularly copper) coherent with µG-linked bone phenotypes and copper’s multifaceted role in immunity and viral life cycles (Man et al., 2022; Albalawi et al., 2024) (Fig. 4B). Several strongly regulated single-stress genes reverted toward baseline in the combination (Fig. S3C): *fipr-4, fipr-7*, and *fipr-10* (defense and cuticle) (Kamal et al., 2022), *grd-17* (hedgehog-like signaling implicated in host–microbe interactions) (Zárate-Potes et al., 2022), and IPR components *B0507.7* and *B0507.8*. Using |log_2_*FC*| > 1, over-expressed genes showing a significant interaction between gravity intensity and infection status were enriched for sphingomyelin biosynthesis and medium-chain fatty-acid catabolism, alongside cellular detoxification; interaction-down genes emphasized cuticle and collagen, a µG hallmark (Çelen et al., 2023), complicated by the antiviral roles of collagens in OrV (Zhou et al., 2024) (Fig. S3D).

**Fig 4.**
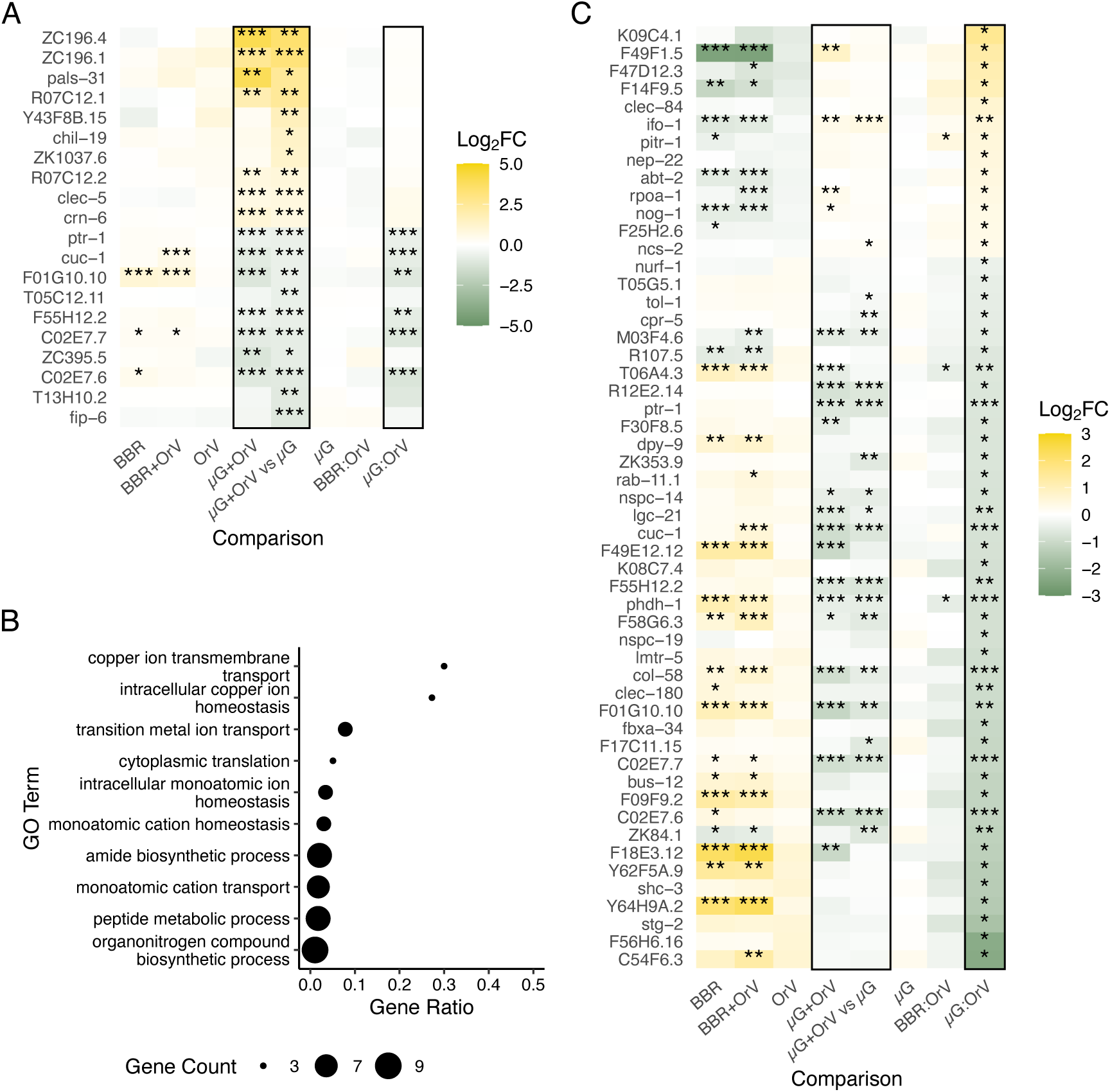
(A) Panel of most significantly upregulated or downregulated genes exclusively under µG conditions after OrV inoculation (14 hpi), compared to non-infected nematodes in this condition. Log_2_*FC* values across all analyzed conditions and interaction effects are shown. Unless otherwise stated, all comparisons were made relative to non-infected nematodes maintained under standard conditions (Std). Asterisks indicate adjusted significance levels (\**P* < 0.05, \*\**P* < 0.01, \*\*\**P* < 0.001). (B) GO enrichment analysis of top biological processes based on exclusively and significantly (adjusted *P* < 0.01) downregulated genes from the comparison in (A). (C) Panel of exclusively and significantly upregulated and downregulated genes for the µG × OrV interaction, displayed as in (A).

After removing genes significantly involved in this interaction, the remaining ones were 13 upregulated and 40 downregulated. (Fig. 4C), with no significant enrichment in any biological process. Most of these exclusive genes exhibit a suppressive combined response that emerges only under dual exposure. The most strongly suppressed gene, *C54F6.3*, a chondroitin 4-sulfotransferase (Mizuguchi et al. 2009), along with collagens and C-type lectins (GO:0005615 (extracellular space), odds ratio = 11.262, *P* = 0.017), could reveal stress combination heightens cellular vulnerability to oxidative or inflammatory damage weakening extracellular matrix-mediated protection. Together with the reduced overall DEGs under µG + OrV, these data reinforce transcriptional attenuation and resource conservation under multi-stress exposure.

### 3.5. Viral phenotypes under stress: replication state, sequence diversity, and nsVG architecture

To assess whether transcriptional antagonism translated to viral fitness consequences, we quantified viral RNA accumulation (viral load) and characterized viral population genomes. Viral load (quantified based on viral reads on the RNA-seq data) decreased under BBR and dropped further under µG, a decline driven largely by RNA2, thereby reducing the RNA2/RNA1 ratio and suggesting infections were earlier in their replication program despite identical sampling windows (Fig. 5A - 5B) (Castiglioni et al., 2024).

**Fig. 5.**
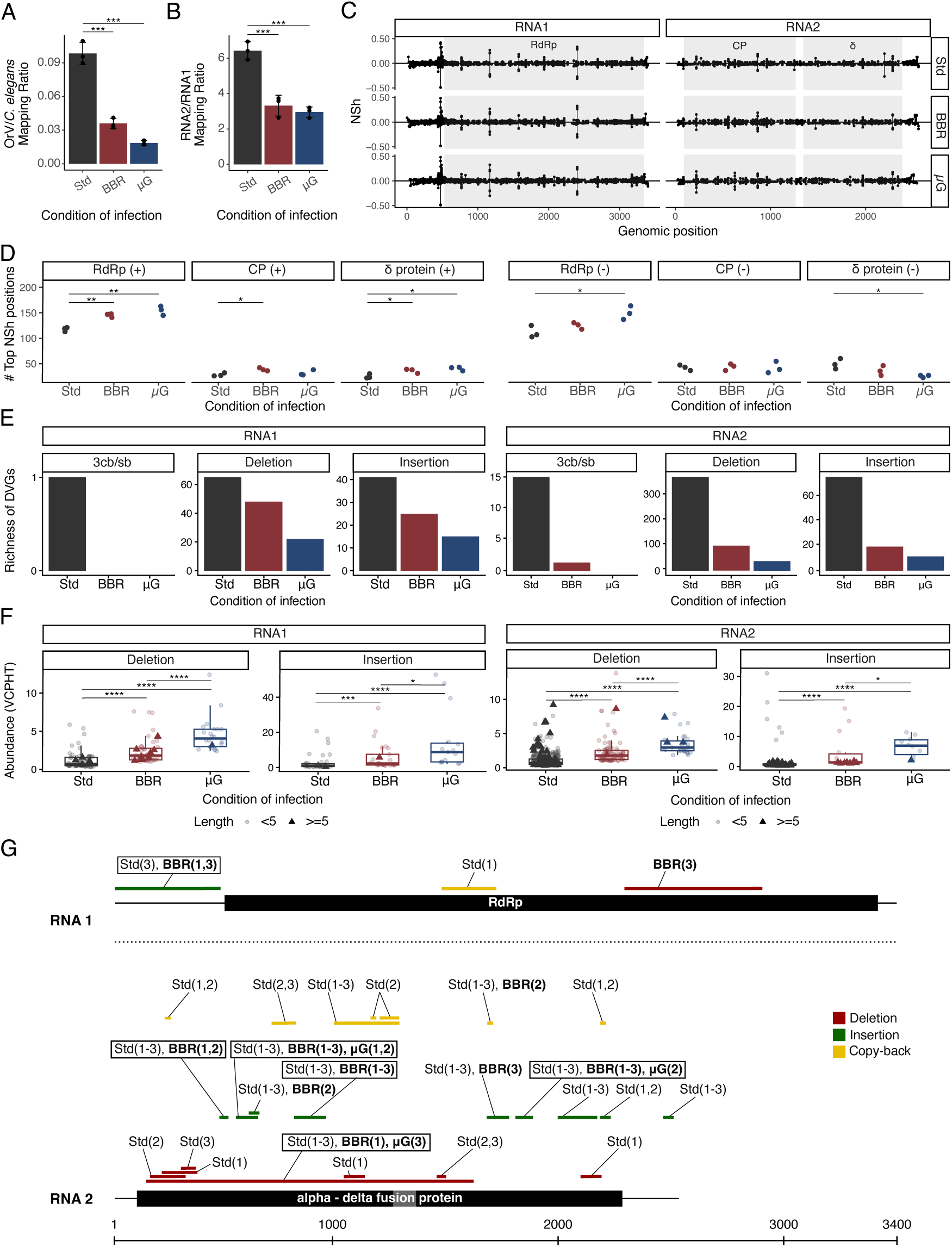
(A) Bar plot showing OrV-to-*C. elegans* mapping ratios across infection conditions, presented as mean ±1 SD. Ratios significantly differed among the three experimental conditions (Welch robust ANOVA: *F*_2,3.145_ = 85.794, *P* = 0.002, η^2^ = 0.977), with differences being due to the reduction of the ratio under the two stresses compared to the standard conditions (Bonferroni *post hoc* test, *P* < 0.001). (B) Bar plot illustrating RNA2/RNA1 mapping ratios across the three infection conditions, with mean ±1 SD. Ratios significantly differed among the three experimental conditions (Welch robust ANOVA: *F*_2,3.638_ = 43.476, *P* = 0.003, η^2^ = 0.939), with differences being due to the reduction of the ratio under the two stresses compared to the standard conditions (Bonferroni *post hoc* test, *P* < 0.001). (C) Normalized Shannon entropy (*H_n_*) values along viral genome sequences; coding sequences (CDS) are indicated by shaded regions. (D) Number of genomic positions within the top 5% most variable sites, with *H_n_* values shown for positive and negative strands. Statistical significance was assessed using a *t*-test; asterisks indicate significance (\**P* < 0.05, **\*\****P* < 0.01). (E) Richness of nonstandard viral genomes (nsVGs), including copy-backs, insertions, and deletions, detected by DVGfinder metasearch algorithm (option ViReMa). (F) Abundance of nsVGs expressed as counts per 100,000 viral reads (VCPHT), detected by DVGfinder (with ViReMa). Statistical significance was evaluated using a Wilcoxon test; asterisks denote significance (\**P* < 0.05, \*\**P* < 0.01, \*\*\**P* < 0.001, \*\*\*\**P* < 0.0001). (G) Consensus nsVGs detected across samples by DVGfinder in consensus mode, indicating the specific samples where events were identified. nsVGs are color-coded by type, and boxed regions highlight those detected in at least five samples and under two or more conditions.

Normalized Shannon entropy profiles were broadly similar across conditions, but the count of high-entropy positions (top 5%) shifted by strand and coding region (Fig. 5C - 5D). On the positive strand, high-entropy sites increased under BBR and µG within RdRp and δ (and CP under BBR); on the negative strand, RdRp also increased under µG, although the overall number of high-entropy sites decreased under µG. Given that RdRp precedes δ and CP in expression (Castiglioni et al., 2024), these data support a replication-timing mechanism modulated by environmental stress. Mechanistically, elevated ROS in stressed hosts may influence viral diversity by introducing RNA lesions that perturb polymerase extension (Akaike & Maeda, 2000; Akagawa et al., 2025), while certain RNA viruses can exploit oxidative cues to tune capping/replication timing (Gullberg et al., 2015).

Non-standard viral genomes (nsVGs) added a structural dimension. With stringent filtering, richness tracked load (Fig. 5E), yet after normalizing by viral reads (VCPHT), insertions and deletions were more abundant under BBR and further increased under µG (Fig. 5F), indicating greater structural variation per unit viral RNA when replication is constrained. High-confidence, conserved nsVGs included duplications in RNA1 (5’ UTR, capsid) and a recurrent RNA2 deletion spanning CP into δ (Fig. 5G), highlighting CP as a hotspot for quasispecies diversification. A BBR-specific > 5-nt RNA1 deletion (absent elsewhere despite higher standard condition loads) reached up to 6.95 VCPHT, consistent with a BBR muon flux host environment that stabilizes specific nsVGs or transient defective-RNA1 compositions.

### 3.6. A working model for non-additive stress integration

Across datasets we observe (*i*) acclimation, (*ii*) antagonism between abiotic stress and infection responses, and (*iii*) proteostasis**-** and resource-centric reprogramming. These patterns converge on a model where chronic environmental stress primes transcriptional circuits that interfere with acute infection responses. We propose that multi-stress exposure activates compensatory circuits (*e.g*., IPR modulation, PERK/ER-stress signaling, redox management, and lipid rerouting) that attenuate hallmark infection cascades while maintaining essential defenses and metabolic balance (Reddy et al., 2017; Kim et al., 2020; Brown et al., 2021; Zheng et al., 2022; Son et al., 2024). The repeated involvement of lipid and collagen/cuticle modules points to barrier function and membrane remodeling as levers through which space-analog stress influences both host resilience and viral replication (Van Voorhies et al., 2020; Çelen et al., 2023; Zhou et al., 2024).

### 3.7. Limitations of the study

The combined µG + BBR + OrV condition could not be assayed due to logistical constraints. The complex logistics of these experiments also limited the feasibility of testing multiple genetic backgrounds and time points: genetic dissection using mutant strains (*e.g*., *sbp-1* and *mdt-15*), time-resolved multi-omics (*i.e.*, transcriptome, proteome, lipidome), and longitudinal nsVG tracking to test causality remain priorities for future work. Mapping conserved modules (*e.g*., PERK signaling, copper homeostasis, collagen/extracellular matrix) in mammalian systems will be essential to anticipate immune modulation and viral reactivation risk during space exploration missions (Atri & Melott, 2011, 2014; Mehta et al., 2017; Tanaka et al., 2017; Restier-Verlet et al., 2021; Wu et al., 2024) and to design effective countermeasures.

## 4. Concluding remarks

Acclimation to simulated microgravity and reduced muon flux dampens the canonical OrV infection program and reconfigures host physiology, yielding antagonistic (non-additive) outcomes across gene modules and viral phenotypes. On the host side, we observe selective attenuation of IPR modules, redistribution of lipid and metal (copper) homeostasis, and recurrent impacts on collagen/cuticle biology. On the virus side, we find reduced load, lower RNA2/RNA1 ratios, strand- and CDS-specific entropy shifts, and increased per-read structural variation, with the capsid region emerging as a mutational hotspot. These findings argue for stress-primed transcriptional attenuation and resource allocation as strategies to limit proteotoxic burden under multi-stress exposure while reshaping replication timing and genome plasticity.

## Data availability

RNA-seq data were deposited at the European Nucleotide Archive (ENA) at EMBL-EBI under accession number PRJEB102402 (https://www.ebi.ac.uk/ena/browser/view/PRJEB102402). All the R code generated for this study are accessible in Zenodo at https://doi.org/10.5281/zenodo.17607908.

## CRediT authorship contribution statement

**Ana Villena-Giménez.**: conceptualization, data curation, formal analysis, investigation, methodology, validation, writing-original draft, writing-review and editing; **Esmeralda G. Legarda**.: conceptualization, data curation, formal analysis, investigation, methodology, validation, visualization, writing-original draft, writing-review and editing; **Rubén González**: conceptualization, supervision, writing-review and editing; **Victoria G. Castiglioni**: conceptualization, investigation, methodology, supervision, validation, writing-review and editing; **Santiago F. Elena**: conceptualization, data curation, formal analysis, funding acquisition, project administration, supervision, writing-original draft, writing-review and editing.

## Acknowledgements

We thank Francisca de la Iglesia (I2SysBio) and Rebecca Hernández-Antolín (LSC) for excellent technical support. The help of María J. Olmo-Uceda (I2SysBio) in bioinformatic analyses is greatly appreciated. We are deeply grateful to Carlos Peña-Garay (LSC) for his commitment and dedication to research in subterranean biology, as well as his constant material and emotional support. Many computations were performed on the HPC cluster Garnatxa at I2SysBio (CSIC-UV).

This study was supported by ESA contract 4000135960/21/NL/GLC/my, grant PID2022-136912NB-I00 funded by MCIN/AEI/10.13039/501100011033 and by “ERDF a way of making Europe”, grant CIPROM/2022/59 funded by Generalitat Valenciana, and grant PB-02-22 from ICTS Laboratorio Subterráneo de Canfranc to S.F.E. V.G.C. was supported by grants FJC2021-047264-I, funded by MCIN/AEI/10.13039/501100011033 and by NextGenerationEU/PRTR, and MSCA 2024-PF-01-101207897, funded by Horizon Europe. E.G.L. was supported by grant FPU21/00410 funded by MCIN/AEI/10.13039/501100011033 and by “ESF invest in your future”. The funders played no role in study design, data collection, analysis and interpretation of data, or the writing of this manuscript.

