## Supplementary figures and images for "Cosmic silence and viral noise: transcriptomic crosstalk in *Caenorhabditis elegans* under simulated space conditions"

### Fig. S1

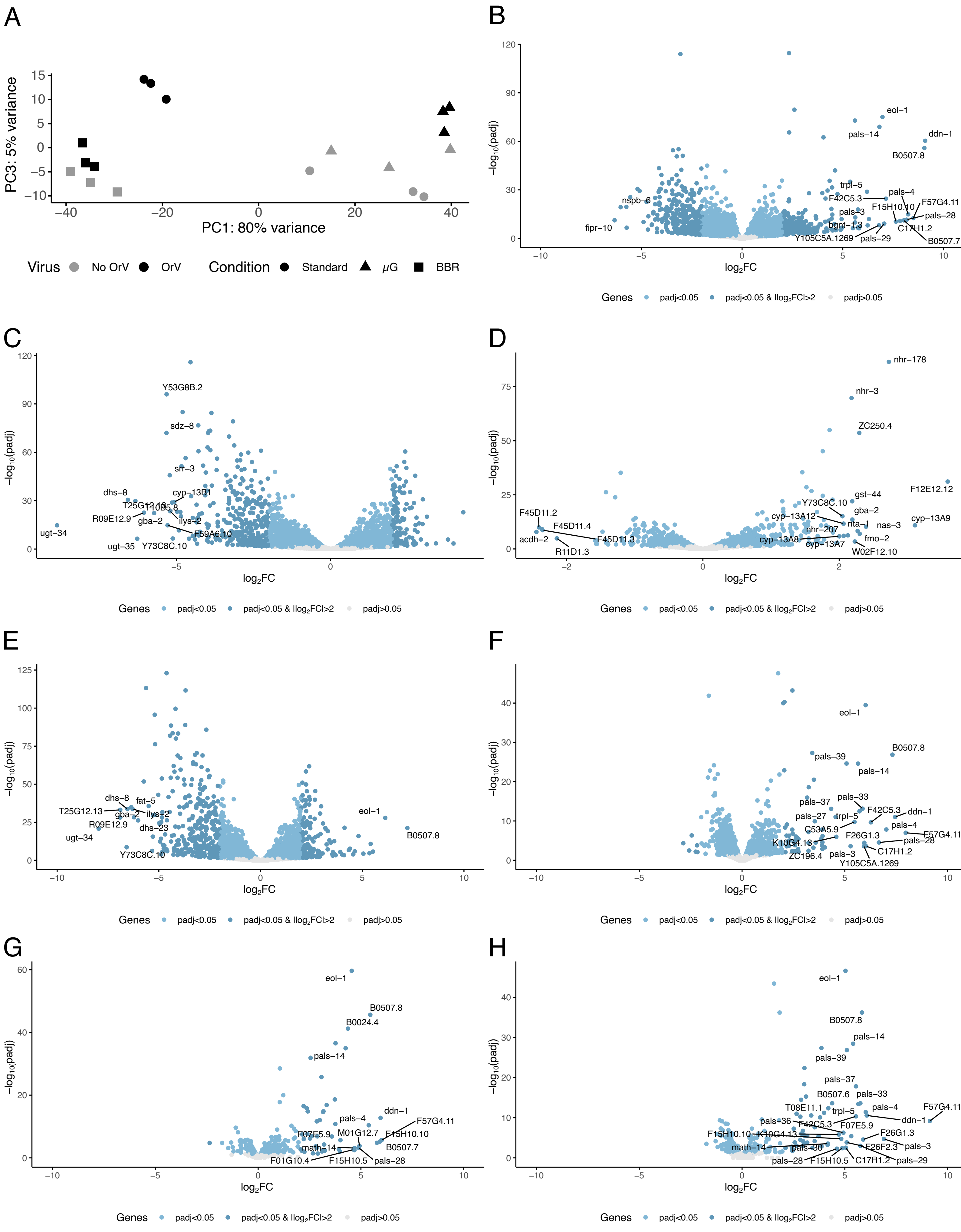

### Fig. S2

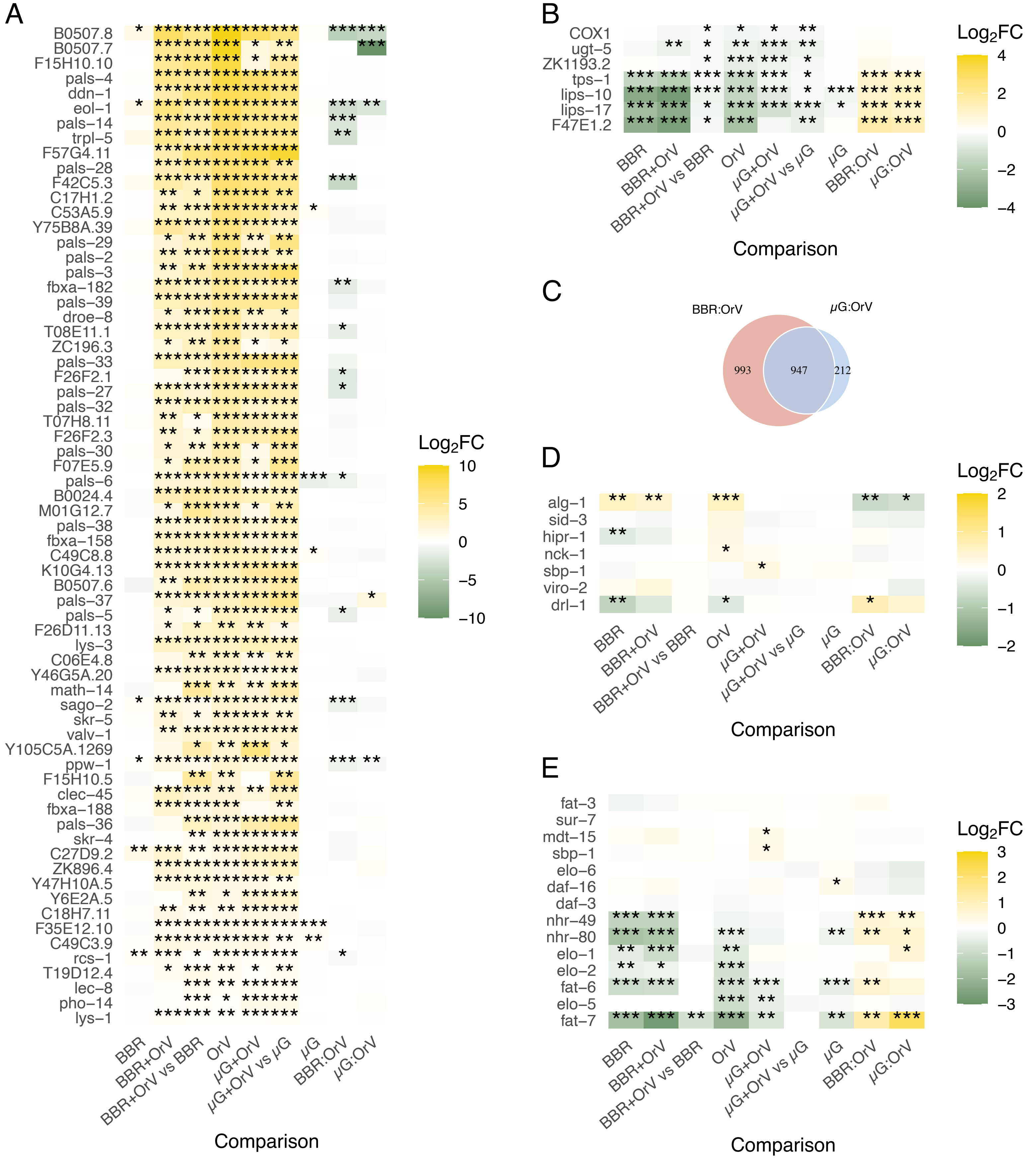

### Fig. S3

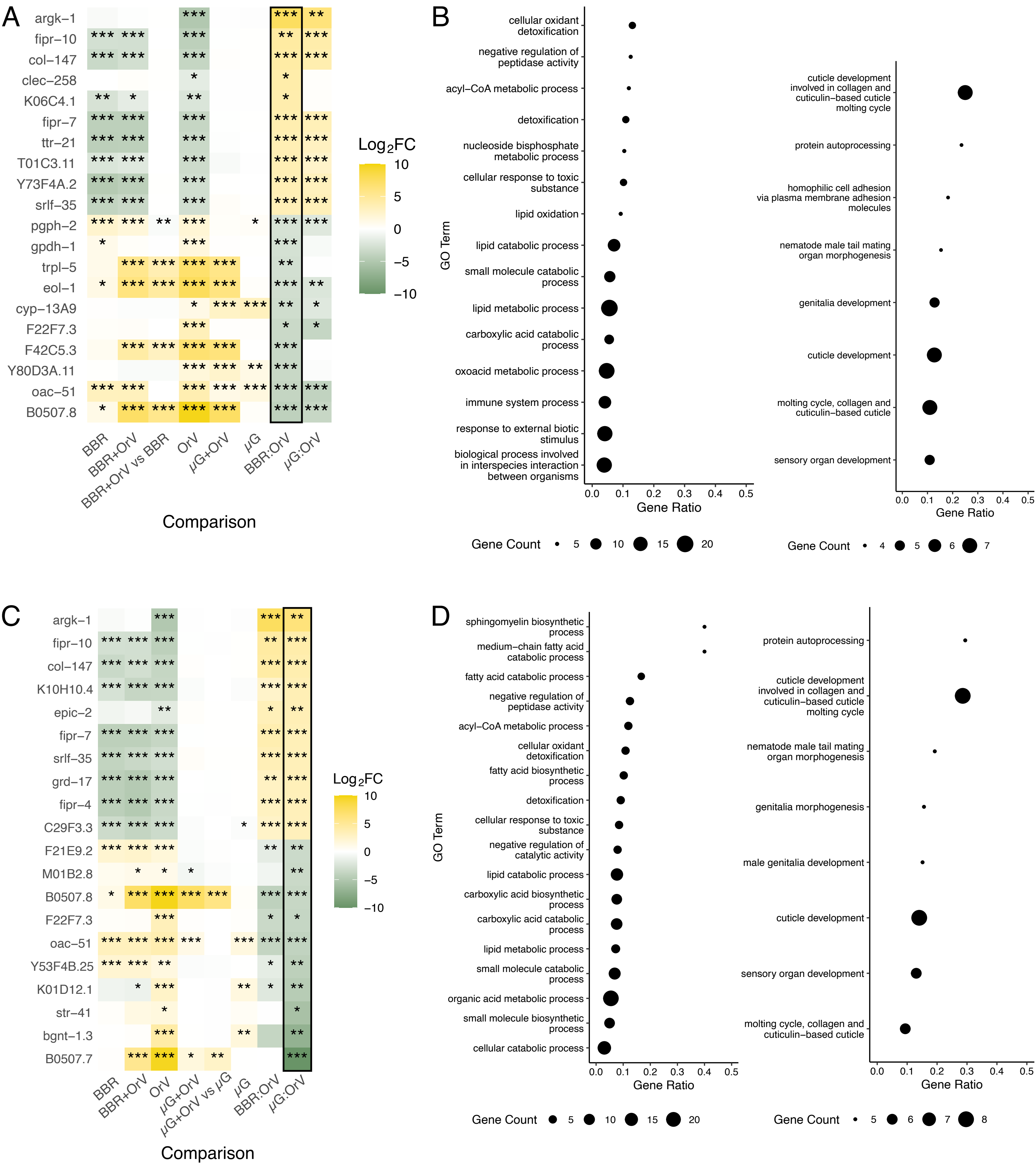
